# Revealing the diversity of light-harvesting antenna of in vivo photosystem I

**DOI:** 10.1101/2025.01.02.631149

**Authors:** Xianjun Zhang, Rin Taniguchi, Shen Ye, Yutaka Shibata

**Affiliations:** Department of Chemistry, Graduate School of Sciences, Tohoku University, Sendai 980-8578, Japan; Division for Interdisciplinary Advanced Research and Education, Tohoku University, Sendai 980-8578, Japan

**Keywords:** photosynthesis, *Chlamydomonas*, antenna system, excitation spectrum, cryogenic optical microscope

## Abstract

The photosynthetic reaction is driven by the cooperation of two light-excited pigment-protein supercomplexes: photosystem II (PSII) and photosystem I (PSI). The efficiency of the excitation of the two PSs relies on the exquisite organization of their light-harvesting antenna under environmental fluctuations. However, since the antenna-protein composition within cells remains elusive, the in vivo events arising from antenna variations cannot be accurately explored. Here, we implemented the single-pixel excitation-emission spectroscopy of *Chlamydomonas* cells under 80 K using a cryogenic optical microscope. The antenna variations of in vivo PSI can be exclusively evaluated via this low-temperature spectro-imaging method. The simultaneous acquisition of two types of fluorescence spectra enables the analysis of the intracellular correlation between the PSII/PSI intensity ratio and the chlorophyll-*b*/*a* (Chl-*b*/*a*) concentration ratio. We found that the Chl-*b*/*a* ratio hardly correlated with the PSII/PSI intensity ratio in most cases, suggesting that the in vivo PSI intensity ratio reflects the relative PSI stoichiometry rather than their antenna sizes. More importantly, the analysis of the PSI antenna-related Chl-*b* concentration within cells reveals a mega-antenna system that is much larger than the antenna sizes of the PSI supercomplexes whose structures have been resolved so far. Such PSI megacomplexes tended to be enriched in the region surrounding the pyrenoids of *Chlamydomonas* cells. We anticipate the present investigation to be a starting point for directly estimating the arrangements of antenna systems of photosystems at the single-cell scale, which is necessary for a deeper understanding of dynamic in vivo events related to the photosynthetic light-harvesting process.

## Introduction

Photosynthetic organisms underpin almost all life on Earth by efficiently collecting and utilizing solar energy (1, 2). Oxygenic photosynthesis requires two photosynthetic apparatuses—photosystem II (PSII) and photosystem I (PSI). They cooperatively drive the photoinduced electron transport from water to the nicotinamide adenine dinucleotide phosphate (NADP+), an essential substance for the Calvin–Benson cycle (2). Solar energy is initially captured by the robust light-harvesting network established within the antenna system, then the excitation energy is rapidly transferred through the antenna array to the reaction center (RC) of the two photosystems (PSs) (2–4). Photon/electron conversion is achieved with a near unity quantum yield in the RCs under ideal conditions (5).

The key to maintaining the high efficiency of the photosynthesis reaction is the efficient excitation energy transfer (EET) between antenna proteins and the PSs (6–8), as well as the balanced excitation between the two PSs (9, 10). However, both PSs are at risk of being overexcited or experiencing unbalanced excitation under dynamically fluctuating light conditions (11–16). Antenna proteins play a crucial role in maintaining the efficient functioning of photosynthesis by not only fine-tuning its own conformation to alter the EET efficiency (17) but also flexibly migrating between the two PSs to adjust their absorption cross section according to the light condition (18–23). Until now, the static antenna structures of the two PS supercomplexes have been resolved in vitro, owing to the drastic advancement of cryogenic electron microscopy (cryo-EM) (24, 25). Nevertheless, there has been a large gap between our current knowledge of the in vitro structures of supercomplexes and issues regarding the in vivo antenna system. For example, we have not yet fully understood the connectivity between the antenna proteins and two PSs and the inter-protein arrangement within an antenna array. The lack of knowledge about in vivo antenna architecture limits the investigation of antenna modulation–related physiological events in a microscopic space. As a result, elucidating the dynamic structures of antenna systems with greater intactness has been strongly desired.

Although the cryogenic electron tomography technique has the potential to access the configuration of photosynthetic components inside the cell, the cellular sectioning makes the observable area extremely limited (< 200 nm) (26). Recently, the atomic force microscope (AFM) has allowed some advances in the characterization of supercomplexes on isolated thylakoid membranes (27–32). However, it is difficult to observe the antenna array on the natural membrane via AFM observation because this technique requires the membrane to be fixed on a substrate surface. Furthermore, AFM has difficulty identifying antenna proteins that have poor portions protruding from membranes. Here, we notice that the chlorophyll-*a* (Chl-*a*) is embedded in all types of photosynthetic proteins, whereas the chlorophyll-*b* (Chl-*b*) is exclusively associated with the peripheral antenna proteins. Hence, the local Chl-*b*/*a* ratio can be a good indicator of the in vivo antenna size of PSs (22, 33).

In this study, we characterize the in vivo antenna system of PSI supercomplexes within *Chlamydomonas* cells using the unique technology of cryogenic single-pixel excitation-emission spectroscopy. By decomposing the spectroscopic components of the excitation and emission spectrum, the Chl-*b*/*a* ratio and PSII/PSI intensity ratio within a cell can be mapped three-dimensionally. We confirmed that PSI is the dominant origin of the cryogenic excitation spectrum, and so the changes in the in vivo Chl-*b*/*a* ratio can reflect the local antenna-size variations of the PSI supercomplex. Our results uncovered a very high heterogeneity of the in vivo antenna organization of PSI. Correlation analysis showed that the local antenna size has little correlation with the PSII/PSI intensity ratio, indicating that the relative fluorescence intensities of PSII and PSI reflect their in vivo stoichiometry rather than antenna size (Fig. S1) (34, 35). Moreover, the antenna size of PSI at each pixel could be facilely estimated in detail by comparing the experimental Chl-*b*/*a* ratios and the ratios of the calculated dipole strength of Chl-*b*/*a* from some assumed PSI structures. These analyses revealed that the antenna sizes of PSIs in the local domains surrounding the pyrenoids are much larger than that of the largest PSI supercomplexes known thus far (36, 37).

The site-dependent antenna architecture revealed by this study indicates the flexible modulation mechanism of the local light-harvesting capability, which may be relevant to the regulation strategy toward the perturbation of dynamic environmental light. This mechanism is tightly related to the physiological function of photosynthetic organisms. This work opens a way for the intuitive analysis of intracellular heterogeneity of light-harvesting antennae.

## Results and discussion

### Cryogenic Single-pixel Excitation-emission Spectroscopy

Single-pixel excitation-emission spectroscopy is realized by the cryogenic excitation spectral microscope (cryo-ESM) we developed.(38). Briefly, in the excitation spectral mode, a multi-color line-shaped laser beam covering the 635–695 nm range is used as the excitation source (Fig. 1A). A long-pass dichroic mirror (DM) with the cut-off wavelength at 700 nm was used to filter the emission spectra. Since excitation-wavelength scanning and spatial scanning are simultaneously achieved in this mode, the cryo-ESM enables the rapid acquisition of the excitation spectra of each pixel on the image. In the emission spectral mode, the cryo-ESM can acquire the emission spectra of cells by facilely replacing the DM with a half mirror (HM) and setting a 640 nm band-pass filter in the excitation pathway (Fig. 1B). To analyze the spectral components of the same pixel on the excitation and emission spectral images, we must estimate and optimize the spatial shift between reconstructed images (SI Text 1 and Text 2, Figs. S2–S4). Additionally, we found that the profile of the low-temperature excitation spectrum of a single cell was severely distorted when the scanning plane was higher than the center plane of the cell. The deformation of excitation spectra may be caused by the absorbance and the light scattering of the excitation laser by the lower part of the cell (SI Text 3, Figs. S5 and S6). Therefore, to accurately acquire the excitation spectra of single cells, the scanning position on the Z-axis was set just at the cell center plane or planes slightly below it for all cells where the deformation of the excitation spectra was negligible.

**Figure 1.**
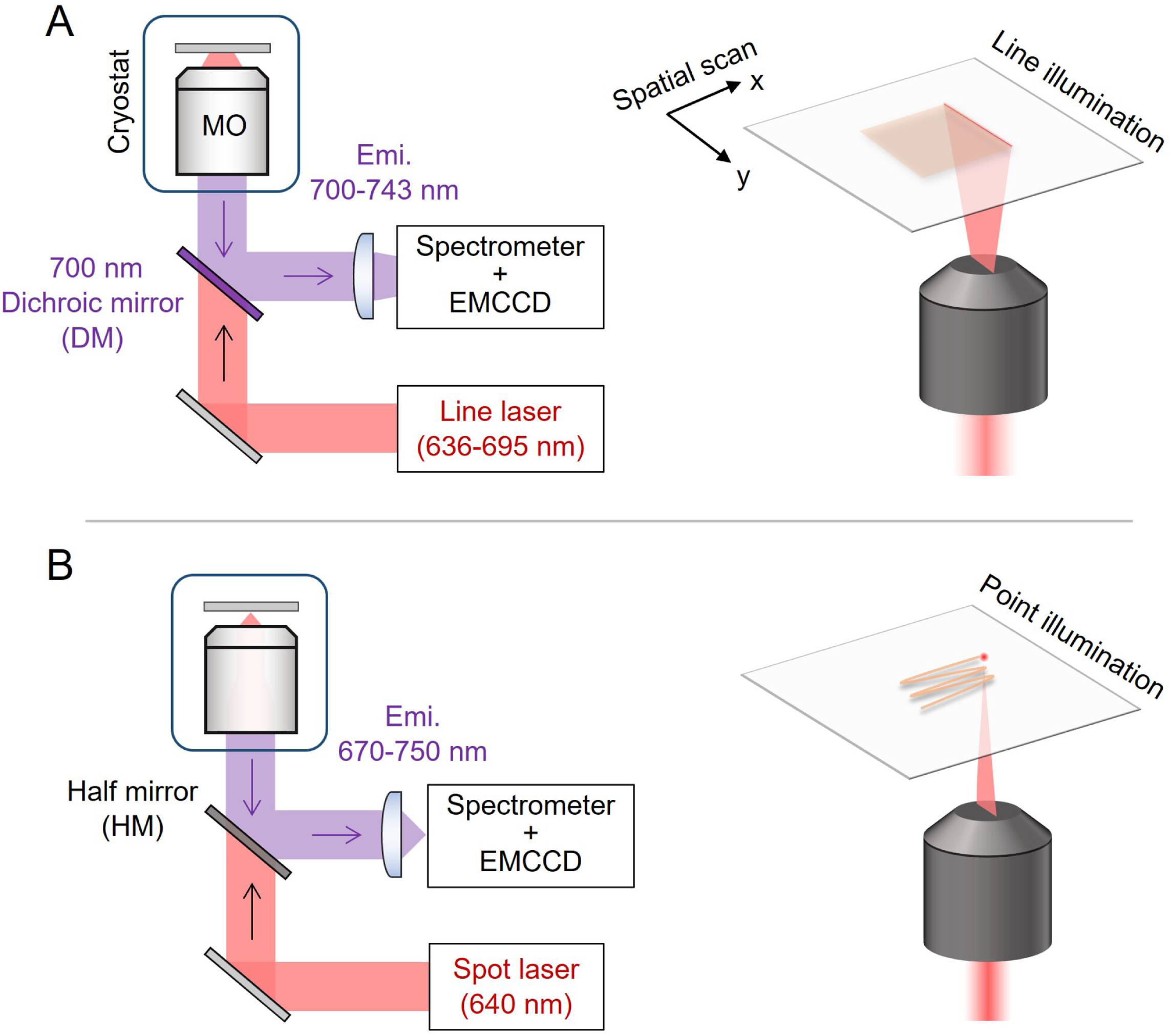
(A) Excitation spectral acquisition mode of the cryo-ESM system. In this mode, a multi-wavelength line laser covering 635–695 nm is collected on the focal plane by a microscope objective (MO). A 700 nm DM filter is used to select the detected 700–743 nm region. The 2D detection mode of the EMCCD camera simultaneously monitors the excitation laser and emission spectrum. The multipoint-excitation and detection configuration uses spatial scanning (x–y direction) to realize the rapid acquisition of the excitation spectra with a high-excitation wavelength resolution of each pixel on the image. (B) Emission spectral acquisition mode of the cryo-ESM system. This mode is equivalent to a traditional confocal laser scanning microscope. The DM is replaced with an HM, and the line laser is changed to a single-wavelength spot-shaped laser by setting a 640 nm band-pass filter in the excitation pathway. The emission spectra with the 670–750 nm wavelength region of each pixel can be obtained.

We performed a cryogenic measurement for 25 *Chlamydomonas* cells at 80 K. Figure 2A–D shows the average excitation and emission spectra of single cells. The PSI fluorescence bands in the 700–730 nm region were significantly enhanced at low temperatures, as shown in panels B and D. We found that the emission spectra of measured cells exhibited characteristics similar to those of cells in state transition (ST). ST is a regulation mechanism that efficiently balances the excitations between PSII and PSI by rapidly shuttling mobile light-harvesting complex (LHC) II between them (11–16). Figure S7 shows the typical emission spectra of cell suspensions undergoing ST induction measured by a conventional fluorometer; they show the spectral characteristics where the PSII and PSI fluorescence emission are enhanced upon the state1 (ST1) and state2 (ST2) inductions, respectively. Although we did not intend to induce the cells into the ST mechanism in the present microscope measurements, we still observed two groups of cells showing different relative intensities of PSII and PSI. Consequently, we categorized all of the measured cells into two types—ST1-like and ST2-like cells—based on the relative emission intensity of PSII and PSI. Figure 2E and F shows the average excitation and emission spectra over the ST1-like (red) and ST2-like (blue) cells, respectively. The ST2-like cells displayed a shape very similar to that of the typical emission spectra of the ST2-induced cells, shown in Fig. S7, whereas the emission spectra of ST1-like cells have relative PSI bands just slightly larger than those of the typical ST1 cell emission spectrum. We found that the number of Chl-*b* components of the ST2-like cells was higher than that of the ST1-like cells (Fig. 2E). It should be noted that the detected emission region was set to 700– 743 nm in the excitation spectral mode. We confirmed that the fluorescence in this region originates predominantly from PSI (SI Text 4). As a result, the enhanced Chl-*b* component on the excitation spectra of the ST2-like cells, as shown in Fig. 2E, reflects the larger antenna size of PSI, i.e., reflects the connection of LHCII to PSI.

**Figure 2.**
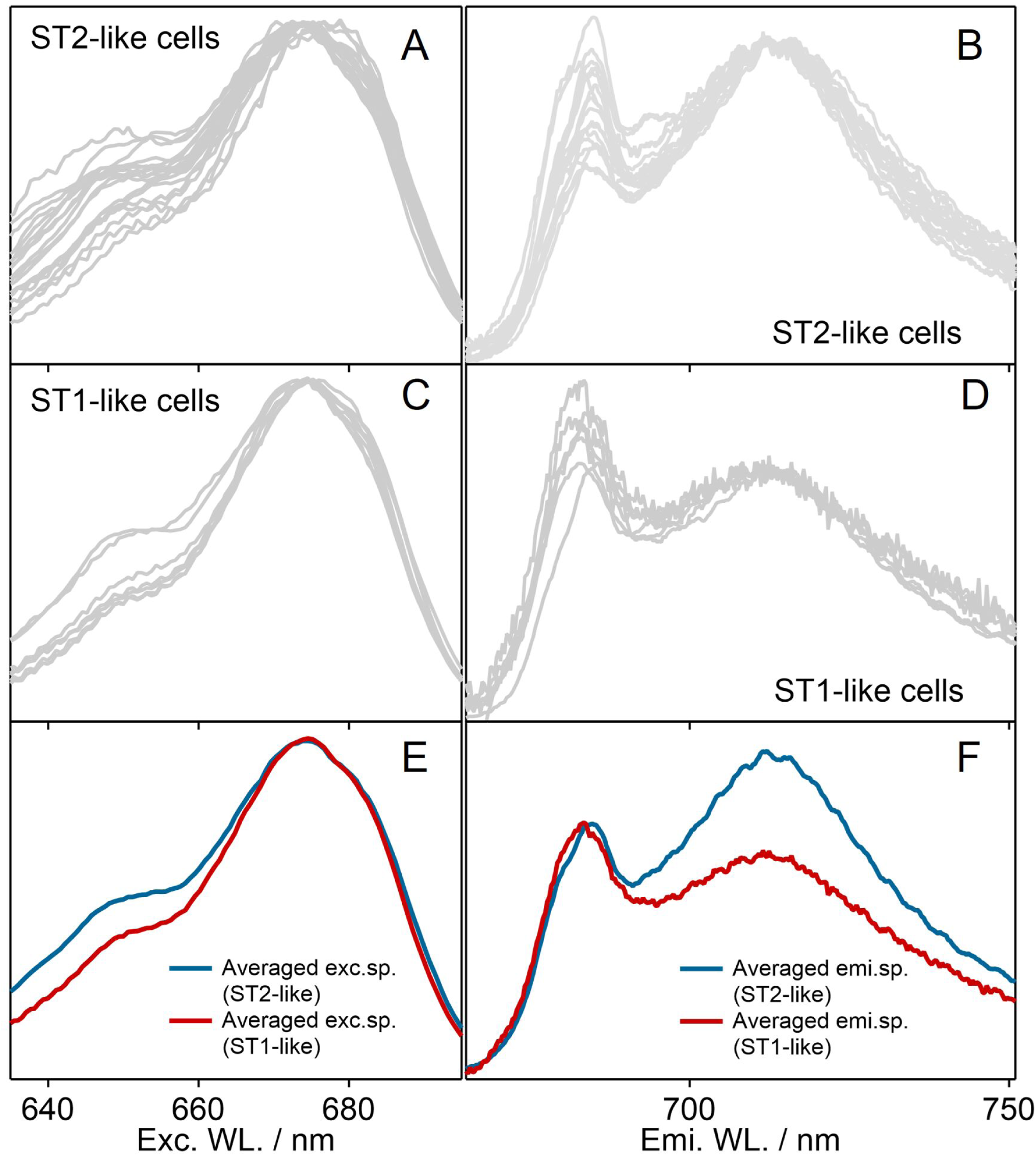
(A–D) The excitation and emission spectra (gray) of 18 ST2-like and seven ST1-like cells at 80 K were measured by cryo-ESM. (E and F) The average excitation and emission spectra of 18 ST2-like cells (blue) and seven ST1-like cells (red).

### Correlation Analysis of the Local PSII/PSI Stoichiometry and Antenna Size

To visualize the in vivo fluorescence components from emission and excitation spectra, we decomposed the emission and excitation spectra by Gaussian fitting and spectral integration, respectively. We fitted the emission spectra to the sum of five-component Gaussian functions. The parameters used in the fitting of ST1-like and ST2-like cells were slightly modified from those of previous works (39). Figure S9 displayed these fitting results in the average emission spectra from two representative cells assigned to ST1-like (panel A) and ST2-like (panel B). Gaussian components with peaks at around 677 nm, 684/692 nm, 710 (713) nm, and 732 nm were assigned to LHCII, PSII_1/PSII_2, PSI, and the vibronic bands of all photosynthetic components, respectively. These components are indicated by green, blue, red, and gray colors, respectively (Fig. S9 and Table S1). Then, emission spectra at all pixels were fitted under a condition where the peak positions and widths were fixed to the values determined by the fitting of the average spectra, and only the amplitudes of each component were allowed to vary. The intensities of the individual Gaussian components of each pixel are given by the products of the width and amplitude of the components. For the decomposition of the excitation spectra into the Chl-*b* and Chl-*a* components, the contributions of the Chl-*b* and Chl-*a* of each pixel were estimated by integrating the intensity over the 640–660 nm and 660–690 nm ranges of the excitation spectra, respectively.

Figure 3A–D shows the image of an analyzed *Chlamydomonas* cell. This cell showed a typical cup-shaped chloroplast. The lobe and base regions of this cell can be easily identified (Fig. 3A; bottom panel). There is no strict definition for the base region of *Chlamydomonas* cells; however, the region surrounding the pyrenoids is usually designated as the base, and the domains other than the base region are designated as the lobe (23, 40–43). Figure 3B and C shows images reconstructed from the excitation and emission spectra, respectively. “Whole” images were obtained from the integrated intensity over the entire wavelength range on the excitation and emission spectra. The “Chl-*b*,” “Chl-*a*,” “PSII,” and “PSI” images were reconstructed by the spectroscopic decomposition described above. The “Chl-*b*/Chl-*a*” and “PSII/PSI” ratio maps were reconstructed by dividing the corresponding images (Fig. 3B and C). In the two ratio maps, the intensity values below a preset threshold were discarded according to the following formula:

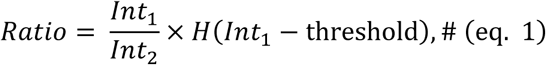

**Figure 3.**
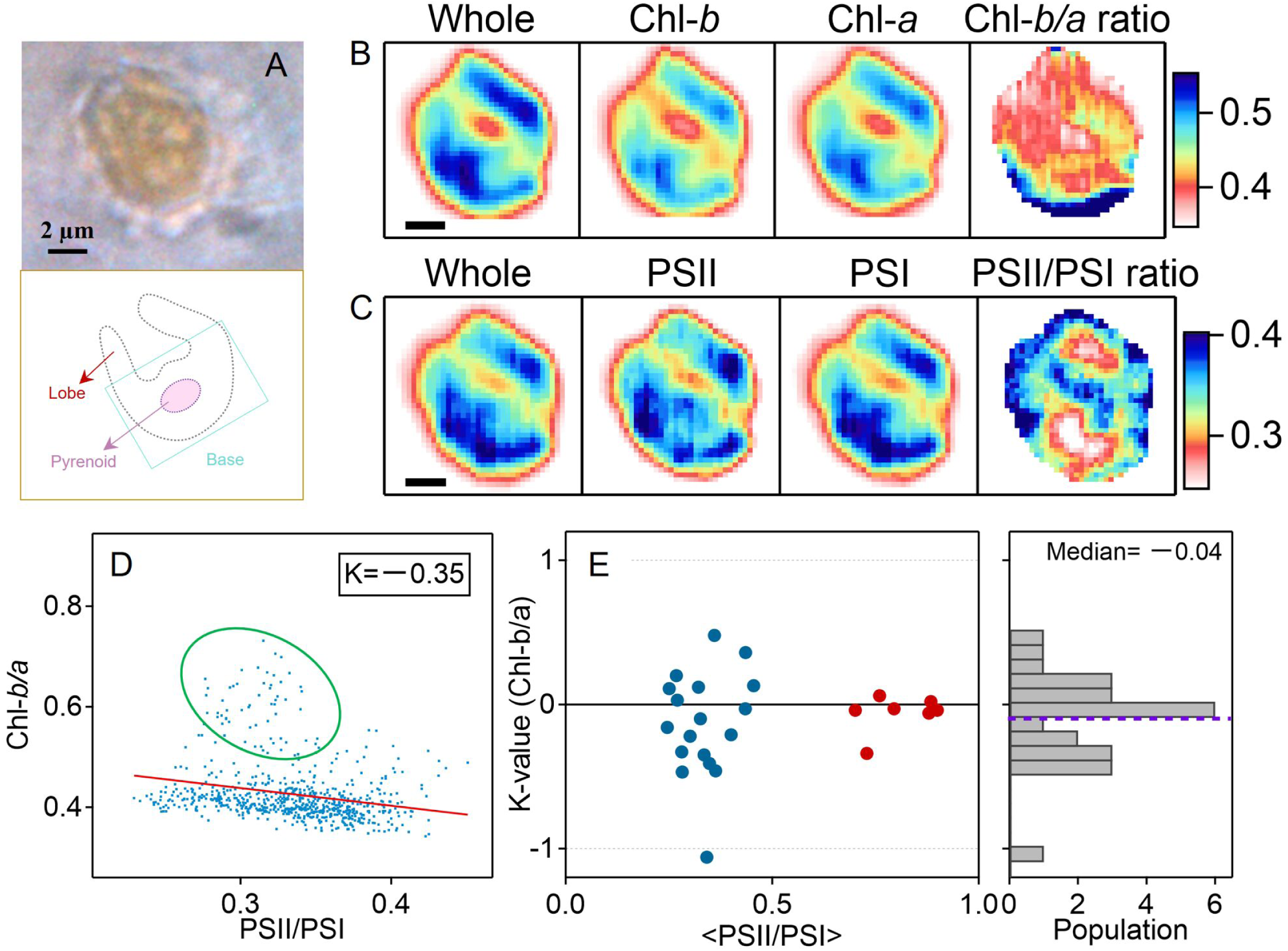
(A) A transmission image of a selected *Chlamydomonas* cell with a typical cup shape. A simple schematic for the profile of this *Chlamydomonas* cell is shown in the bottom panel. (B) Reconstructed images by excitation spectra. The “Whole” image was reconstructed from the intensity signals integrated throughout the entire excitation spectrum. “Chl-*b*” and “Chl-*a*” images were reconstructed from those integrated over the 640–660- and 660–690-nm spectral regions, respectively. The “Chl-*b*/*a*” image is the Chl-*b* ratio map obtained by dividing the “Chl-*b*” images by the “Chl-*a*” images. The scale bar indicates 2 μm. (C) Reconstructed images by emission spectra. The “Whole” image was reconstructed from the intensity signals integrated throughout the entire emission spectrum. “PSII” and “PSI” images were reconstructed by the intensity of individual Gaussian components. The “PSII/PSI” ratio map was obtained by dividing the “PSII” image by the “PSI” image. (D) The correlation plots of the Chl-*b*/*a* vs. the PSII/PSI of this *Chlamydomonas* cell. The green ellipse emphasizes a few larger Chl-*b*/*a* values within this cell. All correlation plots are fitted by the linear function, and the obtained slope is indicated as “K.” (E) The scatter plot of K-values and the PSII/PSI values from 25 cells. Blue and red dots indicate ST 2-like and ST 1-like cells, respectively. The histogram of K-values is shown in the right panel. The median of the histogram is marked by purple dashed lines.

where *H*(*A*) is the Heaviside step function. *Int*_1_ and *Int*_2_ indicate the fluorescence intensity of the corresponding images that are set in the numerator and denominator, respectively.

From the “PSII/PSI” ratio maps, we defined the intracellular regions with a higher and lower PSII/PSI intensity ratio as “PSII-rich” and “PSI-rich” sub-domains, respectively. The low-temperature excitation spectra are very sensitive to variations in the antenna size of PSI, enabling the evaluation of the correlation between the antenna size of PSI and the PSII/PSI intensity ratios. We plot the Chl-*b*/*a* ratios against the PSII/PSI ratios of the same pixels from the two ratio maps (Fig. 3D). The scatter map was fitted to a linear function, and the slope, or K-value, was obtained. K-values intuitively reflect the correlation of the intracellular PSI intensity relative to their antenna size (i.e., Chl-*b*/*a* ratio). If the relative intensity of in vivo PSIs is purely determined by their antenna sizes, the K-value between the PSII/PSI ratios vs. the Chl-*b*/*a* ratios should be equal to −1. On the contrary, a low absolute K-value reflects a situation where the antenna size of PSI is not predominantly correlated to the local relative PSI fluorescence. In this case, the PSII/PSI value is more sensitive to the PSII/PSI stoichiometry than to the antenna size of PSI (Fig. S1A).

We observed a low negative K-value (−0.35) for this cell, indicating a weak colocalization of the Chl-*b* and the relative PSI fluorescence within this cell (Fig. 3D). However, we found that Chl-*b*/*a* ratios in some of the PSI-rich regions (surrounded by the green ellipse in Fig. 3D) exceeded 0.5. If we remove these data points, the estimated K-value will get closer to zero. This result suggests that the local PSII/PSI ratio values of only a few domains are more sensitive to the antenna sizes of PSI than to the PSII/PSI stoichiometries, although such intracellular regions are exceptional (the green ellipses of Figs. 3D and S10). Figure 3E displays the slopes (K-value) vs. the PSII/PSI ratios from all 25 cells measured. The averages of the PSII/PSI ratios are different between these cells, reflecting their ST1-like and ST2-like characteristics. Eighteen and seven cells were categorized as ST2-like and ST1-like types, respectively. The distribution of K-values was narrower for the ST1-like cells than for the ST2-like cells. The right panel in Fig. 3E is the histogram of the K-values from all cells. The median of the distribution is only −0.04. Most of the cells showed a rather small absolute K-value that falls within the range of −0.5 to +0.5, while only an exceptional cell showed a high negative K-value of about −1 (Fig. 3D and Fig. S10). Collectively, considering the two situations shown in Fig. S1 and the colocalization analysis given here, these measurements allow us to conclude that, although a few intracellular areas may imply the inhomogeneous antenna size model shown in Fig. S1B (Fig. S10), the PSII/PSI fluorescence intensity can reflect mainly the PSII/PSI stoichiometry (Fig. S1A).

### Inhomogeneity of Antenna Compositions of PSI

Although it is justifiable to use the slopes of the fitting lines to characterize the in vivo link between the PSII/PSI intensity ratio and the antenna size of PSI, the inhomogeneities of the antenna-complex organizations of PSI have not been fully described only by the K-value, especially for the characteristic intracellular local domains. However, although the 640–660 nm region on the excitation spectrum was preliminarily assigned to the Chl-*b*, this spectral region also covers the vibronic bands of Chl-*a* molecules. Thus, the measured Chl-*b*/*a* ratio cannot be used to accurately estimate the antenna composition of PSI. Figure S11A shows the excitation spectrum of the PSI solution isolated from a thermophilic cyanobacterium, *Thermosynechococcus* (*T.*) *vestitus*, measured by the cryo-ESM. Since cyanobacteria do not contain Chl-*b*, the ratio of the integral intensity in the 640–660 nm to 660–690 nm region on the excitation spectrum of cyanobacterial PSI can be a good reference of the background value of the ratio due to the Chl-*a* vibronic bands. The ratio of the PSI solution of *T. vestitus* was estimated to be 0.312. Therefore, all measured Chl-*b*/*a* ratios for *Chlamydomonas* cells have been corrected by subtracting 0.312, hereafter.

To analyze both intercellular and intracellular diversities in the antenna size of PSI, we plotted the scatter map of the corrected Chl-*b*/*a* ratio vs. the PSII/PSI ratio sampled from 25 *Chlamydomonas* cells (Fig. S11B). The data points in Fig. S11B show a clear negative correlation between the Chl-*b*/*a* ratio and the PSII/PSI ratio. It must be emphasized that this result does not contradict the conclusion given in the previous section based on the K-values falling within the range of −0.5 to +0.5. This apparent contradiction arises because the negative correlation seen in Fig. S11B is mainly due to the intercellular heterogeneity rather than intracellular correlation. These data points seem to be roughly categorized into four groups (I, II, III, and IV). The I and IV groups mainly come from ST1-like cells showing high PSII/PSI ratios, and the II and III groups mainly come from the ST2-like cells with low PSII/PSI values. Figure S11C plotted the medians of the Chl-*b*/*a* ratios against the average of the PSII/PSI ratios of the 25 individual cells. The error bar in Fig. S11C indicates the standard deviation (SD) within each single cell. The SD value quantified the variations in the antenna size of PSI within a cell.

A few biochemical measurement-based studies have suggested that the in vivo synthesis rate of photosynthetic proteins in *Chlamydomonas* cells depends on the period of the cell cycle (44, 45). Therefore, it is expected that the heterogeneity of the antenna-complex composition within a single cell may decrease as the cell ages toward the later period of the cell cycle. In our microscopy experiments, we tended to select larger cells for observation, resulting in preferential observations of cells in a later stage of the cell cycle. Although these cells did not show much difference in cell size, the heterogeneities in the contents of photosynthetic proteins caused by the differences in the synthesis rate cannot be completely excluded. Consequently, the intercellular heterogeneities of the antenna size of PSI may reflect differences in the relative content of PSI and LHCII within each cell (Fig. S11C). We also found that variations of the PSI antenna size in ST2-like cells are larger than those in ST1-like cells. This is reasonable because the organizations of the PSI antenna are more diverse due to the involvement of the additional mobile LHCII during the ST1 to ST2 transition (25, 37).

To further refine the classification of the data points shown in Fig. S11B, we performed a clustering analysis on all pixels by the Gaussian mixture model (GMM)-based machine learning method. The GMM analysis aims to accurately categorize all pixels into certain clusters. We estimated the appropriate number (N=6) of clusters for the present data points based on the Bayesian information criterion (BIC) (SI Text 5). The classified pixels are shown in Fig. 4A. Data points belonging to different clusters are shown with different colors. Each cluster is numbered from C1 to C6, as shown. The C1/C2/C3 and C4/C5/C6 clusters mainly originated from ST2-like and ST1-like cells, respectively. Figure 4B shows the plot of the average of Chl-*b*/*a* ratios against the average of the PSII/PSI ratios of each cluster. The error bars indicate the SDs of the Chl-*b*/*a* ratios and the PSII/PSI ratios of the individual clusters. In 13 of 25 cells, the pixels were classified mainly into only one cluster. In the remaining cells, the intracellular pixels were classified into multiple clusters, strongly reflecting an intracellular inhomogeneity of the antenna size of PSI (Fig. S13).

**Figure 4.**
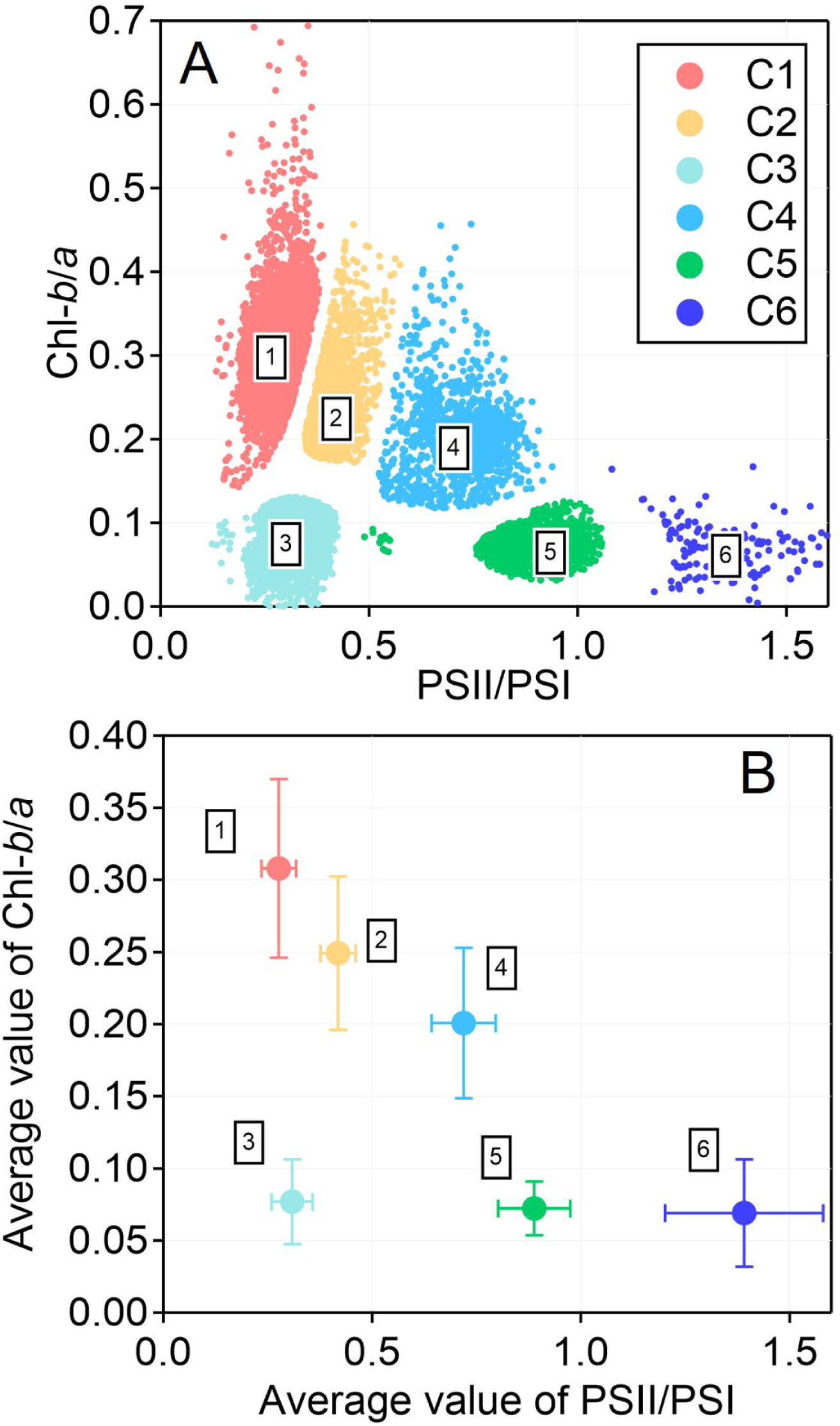
(A) The scatter map of the Chl-*b*/*a* ratio vs. the PSII/PSI ratio composed of all of the pixels from the images of 25 cells. All data was classified into six clusters by GMM analysis. (B) The averages and standard deviations of the Chl-*b*/*a* and the PSII/PSI ratios on each cluster.

### Visualization of the in vivo Distributions of Antenna Composition of PSI

To obtain better insight into the intracellular heterogeneity of the antenna organization of PSI within individual cells, we visualized the intracellular distribution of each cluster. In Fig. 5, we display five selected ST2-like cells that exhibited typical cup-shaped fluorescence distributions. The left and right panels in Fig. 5A-E indicate the reconstructed excitation spectral images and their cluster maps, respectively. For Cell_8, the minor C4 components are visible, which are uniformly distributed in a few pixels shown in blue (Fig. 5A). Since this C4 component is negligible for ST2-like cells, we pay more attention to the C1/C2/C3 clusters. We found that the C1 and C2 components are dominantly distributed in the base region for almost all cells except for Cell_11, in which C1 components extended to the lobe. The C1/C2 clusters exhibited higher heterogeneities of the distribution of Chl-*b*/*a* ratios, indicating a large fluctuation in the antenna organization of PSI. More significantly, the C3 components are predominantly distributed in the lobe region for these cells. Owing to the C3 cluster indicating the lowest Chl-*b*/*a* ratio group, i.e., the smallest antenna size of PSI here, these cluster maps strongly suggest a tendency that the antenna size of PSI in the lobe region is smaller than that in the base region in *Chlamydomonas* cells. On the other side, since the SD of Chl-*b*/*a* ratios in the C3 cluster was relatively low, these observations also reflect that the diversity of antenna size of PSI was lower in the lobe region.

**Figure 5.**
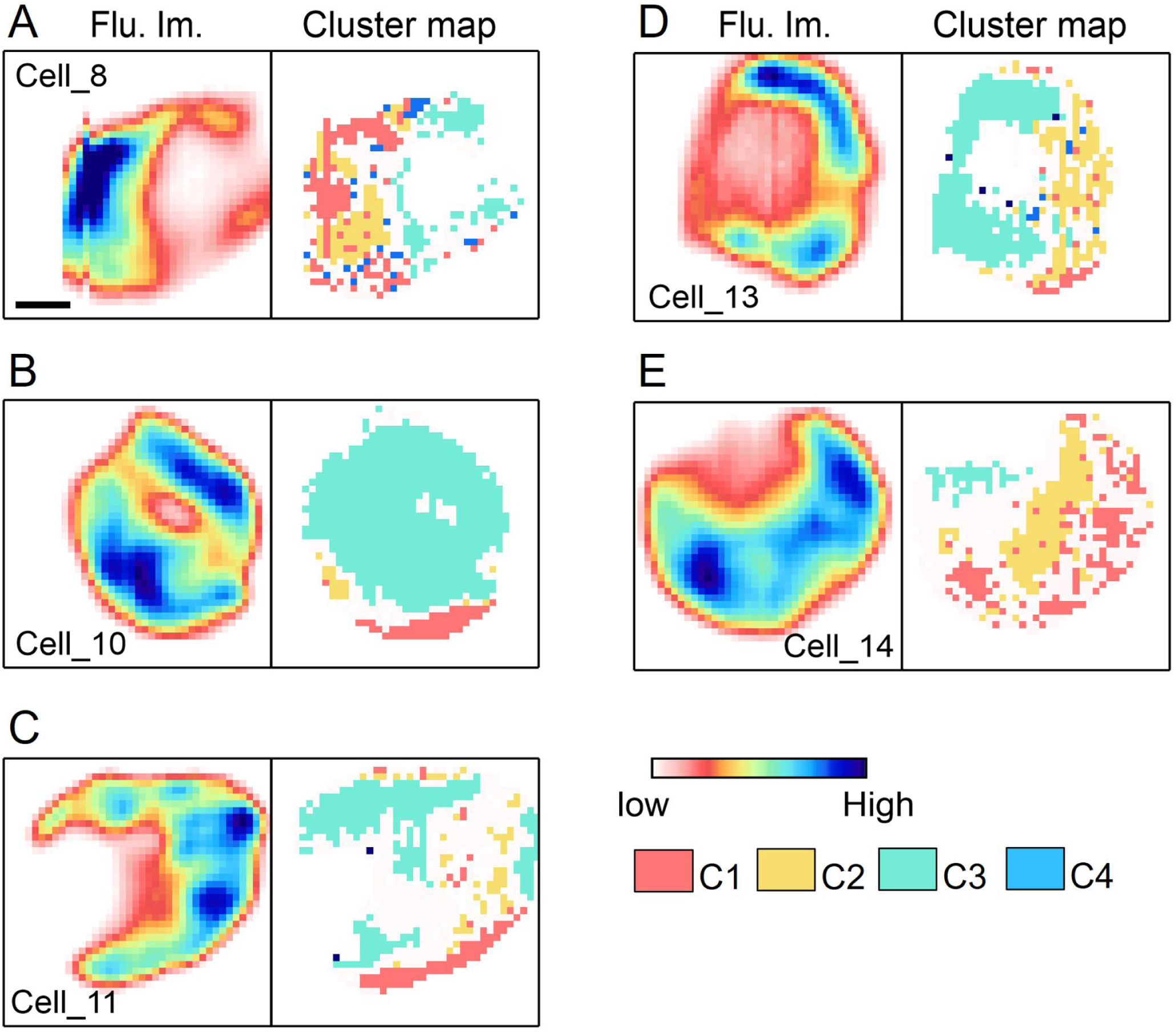
(A–E) Five selected *Chlamydomonas* cells. Their excitation spectral images and cluster maps are shown in the left and right panels, respectively. Serial numbers of each cell correspond to those shown in Figs. S10 and S13.

### Estimation of the in vivo Antenna Composition of PSI

The structures of many photosynthetic supercomplexes have been determined in detail via cryo-EM (36, 37, 46). Until now, two main structures of PSI supercomplexes of *Chlamydomonas* cells, PSI–LHCI and PSI–LHCI–LHCII, have been reported (25, 36, 37). The PSI–LHCI–LHCII supercomplex is formed only after the involvement of mobile LHCII upon ST2 induction. These advancements have provided accessibility to information about the compositions and arrangements of the Chls embedded in these supercomplexes; however, the in vivo antenna compositions have remained largely unknown.

On the basis of the structural information and the assumption of uniform energy fed from Chls-*a* and Chls-*b* to the pigments emitting the fluorescence at 715 nm, it is possible to estimate the area ratios of the Chl-*b* and Chl-*a* components on the excitation spectra of the PSI supercomplexes with different antenna sizes. Owing to the experimental Chl-*b*/*a* ratio obtained by integrating the main spectral regions of Chl-*b* and Chl-*a* on the low-temperature excitation spectra, we can provide a method that simulates the relative spectral areas of the Chl-*b* and Chl-*a* components on the excitation spectrum of a supercomplex. This calculation is based on the squared transition dipole moment 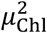 of each pigment multiplied by its number bound to the supercomplex. Since the value of 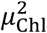 is proportional to the absorption band area of the pigment, 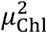 reflects the spectral areas better than the molar extinction coefficients (47). We used a series of putative model PSI supercomplexes, in which different numbers of mobile LHCII trimers were assumed to be bound as the peripheral antenna (Table S2). The Chl-*b*/*a* ratios in these model PSI structures can be calculated according to eq. 2:

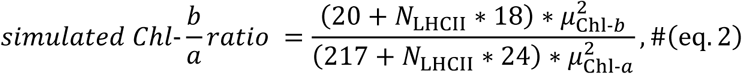

where *N*_LHCII_ is the number of LHCII trimers, 217 and 20 are the quantities of the Chl-*a* and Chl-*b* molecules in the PSI–LHCI supercomplex (Table S2), respectively, and 18 and 24 indicate the numbers of Chl-*b* and Chl-*a* bound to an LHCII trimer, respectively. The values of 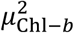 and 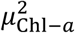 were assumed to be 14.7 D^2^ for Chl-*b* and 21.0 D^2^ for Chl-*a* molecules, respectively. These two values were adopted from the reports by Müh et al. (47, 48). Figure S14 shows the dependence of the simulated Chl-*b*/*a* ratios against the number of additional LHCIIs in the putative PSI supercomplexes. Furthermore, the simulated result (black line) in Fig. S14 was corrected by a factor of 1.27 according to eq. 3:

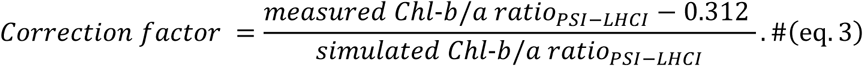

The numerator of eq. 3 was obtained by the measurement of the excitation spectrum of a purified PSI–LHCI supercomplex solution sample with the same setup (inset graph in Fig. S14). Since the supercomplex in this sample is known to bind 20 Chl-*b* and 217 Chl-*a* (see Table S2), the above factor was introduced to correct the Chl-*b*/*a* ratio measured by the cryo-ESM with that estimated according to eq. 2.

We found that the simulated Chl-*b*/*a* ratios were only about 0.082 and 0.18 for the PSI–LHCI and PSI–LHCI–LHCII complexes, respectively (Table S2), in which the value of 0.082 is in good agreement with the average Chl-*b*/*a* ratio (0.077) of the C3 cluster obtained via our microscopy measurements (Fig. 4B). Therefore, the Chl-*b*/*a* ratios of the C3 cluster can be roughly explained as PSI–LHCI complexes. Since the pixels assigned to the C3 cluster were mainly observed in the lobe regions (Fig. 5), the PSI–LHCI complexes may be the main structure of the PSIs localized in the lobe region. Interestingly, the C1 and C2 clusters show higher average Chl-*b*/*a* ratios (0.31 and 0.25). According to the above estimation summarized in Fig. S14, the number of extra LHCIIs bound to PSI–LHCI for the C1 and C2 clusters can be estimated to be six and four, respectively. In some cases, PSIs may be connected to more than ten LHCII trimers (Chl-*b*/*a* ratio > 0.38), especially in C1 clusters. Additionally, the SDs of the Chl-*b*/*a* ratios within the C3 and C1/C2 clusters were about 0.05 and 0.13/0.11 respectively. These results suggest that the PSIs localized in the base region may be energetically associated with a megascale antenna system and exhibit larger variability in antenna compositions (Fig. 4 and Fig. 5).

Another PSI-containing structure, the PSI–PSII megacomplex, was also reported; however, until now, the antenna composition of this structure has not been resolved because of the difficulty of the extraction process (49). Recently, the structure of the PSI–PSII complex was deduced by Kim et al. based on a rough electron microscope observation (50). In this deduced structure, the PSI– LHCI is directly associated with about seven LHCII trimers. To the best of our knowledge, this is the largest antenna size of PSI among the structure-reported PSI supercomplexes. Nevertheless, even if we assume that all of the excitation energy is fed to PSI in this complex, this structure gives a simulated Chl-*b*/*a* ratio of 0.34, which is close to the values of the C1 cluster obtained via cryo-ESM. It should be noted here that the present experiment did not capture evidence for the existence of the PSI–PSII megacomplex. We just utilized the information on the pigment composition of this complex to verify whether the existence of the PSI–PSII complex can explain the large Chl-*b*/*a* ratio observed in the present study. Indeed, it partly explains the experimental Chl-*b*/*a* ratios obtained in the C1 cluster. To confirm the PSI–PSII megacomplex, an experiment demonstrating the direct energy transfer (“spillover” mechanism) from PSII to PSI is necessary (49, 51). This is beyond the scope of the present study. Another study surveyed the relative stoichiometry of LHCII and PSI on the *Chlamydomonas* thylakoids under different culture lights (34). Their measurements showed that the ratios of LHCII/PSI are variable in the range of about 1.15 to 1.55, which are far smaller than the values estimated by our results. Therefore, these experiments suggest a difference between the antenna sizes of in vivo and in vitro PSIs.

### Structure Model of in vivo PSI Supercomplexes

Based on the observations and analysis of the high heterogeneity of antenna proteins for intracellular PSIs, we obtained new insights, as depicted in Fig. 6. The PSI–LHCI supercomplex can be regarded as the main structure of PSI at the lobe, whereas the PSIs at the base may exist in the form of megascale complexes. Such megascale PSIs are energetically associated with about four to six more LHCIIs than are the PSIs at the lobe region (Fig. 6). Two thought-provoking questions can be raised: how do the additional LHCIIs connect to PSI–LHCI, and why do the PSIs with large antenna systems tend to be localized at the base?

**Figure 6.**
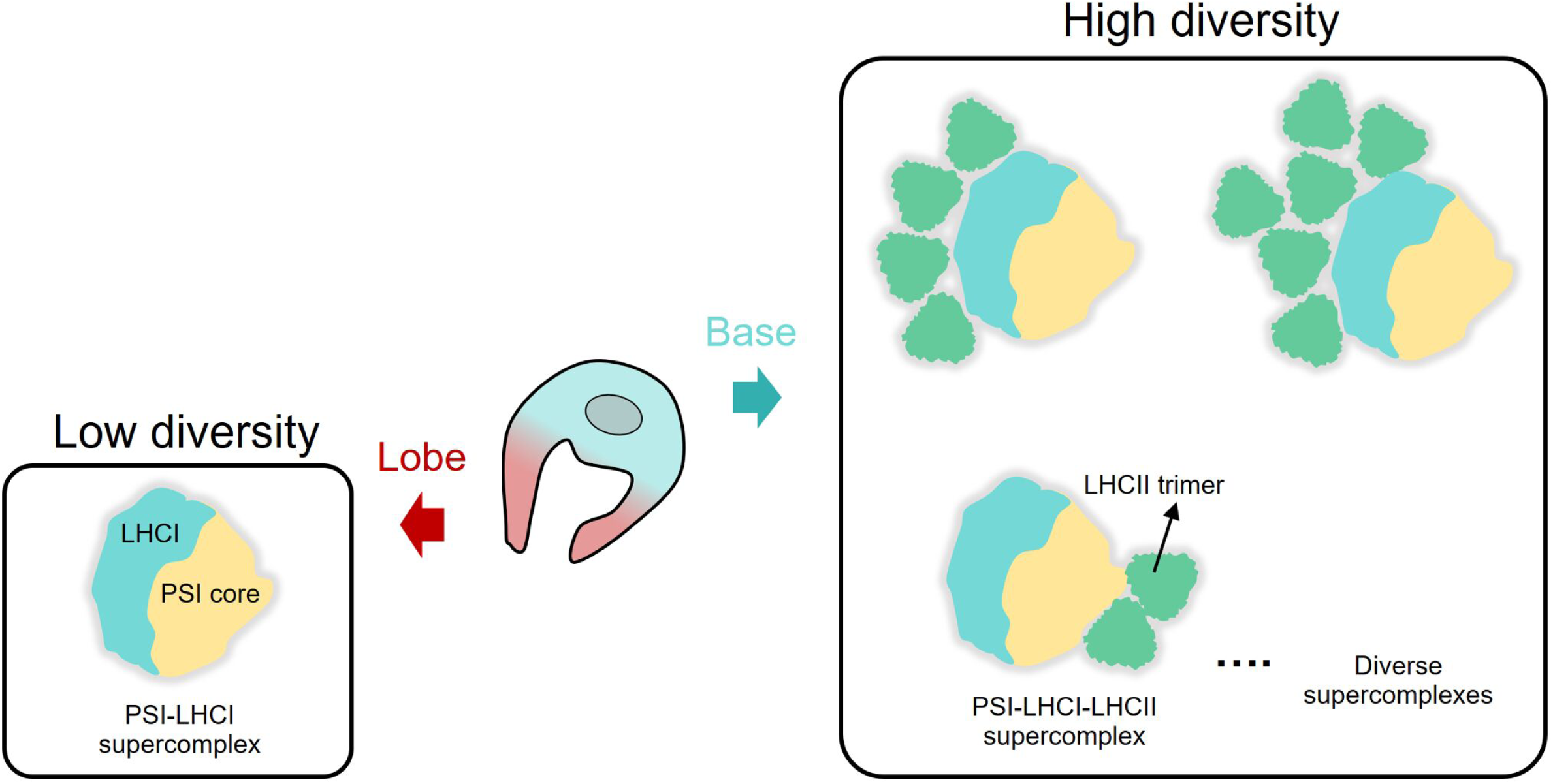
Putative antenna composition of the PSI supercomplex within a *Chlamydomonas* cell. The base and lobe regions of a cell are filled with light blue and light red, respectively. The PSI–LHCI supercomplex (PDB code: 6JO6), PSI–LHCI–LHCII supercomplex (PD code: 7DZ7), and the LHCII trimer (PDB code: 7DZ7) are roughly drawn. The PSI core and the LHCI belt are indicated in yellow and light blue-green, respectively. The C1 and C2 clusters shown in Fig. 4A suggest the two main PSI structures in the base region. Such PSI–LHCIs in the base region additionally bind four to six more LHCII trimers (triangles filled with blue-green).

We can simplify the first question: are the additional LHCIIs directly bound to the PSI core as in the PSI–LHCI–LHCII supercomplex or indirectly via the LHCI belt? A time-resolved fluorescence spectroscopy study has reported that the fluorescence emission of the red Chl at around 715 nm originated from the LHCI belt rather than from the PSI core (52). Therefore, if the extra LHCIIs connect directly to the PSI core, the excitation energy captured by LHCII can hardly reach LHCI due to the long distance and intercept by the PSI core part. In this case, the fluorescence emission from the red Chl will not be enhanced by the additional LHCII, and high heterogeneity in the fluorescence excitation spectra will not be observed. This is obviously inconsistent with the present results. Accordingly, we suggest that the extra LHCII trimers contributing to the increased Chl-*b*/*a* ratio should connect to the LHCI belt side (Fig. 6). An important future work would be to explore the connectivity between LHCIIs and LHCI within the PSI supercomplex in detail.

For the second question, we consider that such exquisite organization of PSI may involve the mechanisms by which the antenna configuration controls light harvesting. In our recent work, using room-temperature ESM, we reported a site-dependent ST mechanism in *Chlamydomonas* cells where the base region shows less susceptibility to ST induction (23). In that study, we tentatively assumed possible situations that are responsible for the observed lack of ST activity in the base region. In the present work, cryo-ESM obtains better spectral contrast for PSII and PSI fluorescence so that the above observation can support one of the possible explanations for the occurrence of inhomogeneous ST: the PSIs in the base region have larger antennae and tend to stay in ST2. Therefore, we also suggest that the PSI–LHCI–LHCII structure may widely exist in the base region (Fig. 6). It is worth noting that the pyrenoid localized at the base is the subcellular organelle responsible for CO_2_ fixation catalyzed by the enzyme Rubisco. Many studies have shown that the efficiency of RuBisCO reacting with the substance CO_2_ is rather low, and CO_2_ is competitively substituted by O_2_ (53–56). Hence, the CO_2_ fixation activity of RuBisCO is drastically diminished under the existence of oxygen. The PSIs with larger antenna sizes also may be of physiological importance because such organizations at the base region may reflect the enhanced efficiency of the cyclic electron transfer (CET) via PSIs. Since the electron transfer in the CET does not require electron flow from PSII, enhanced CET activity in the region surrounding the pyrenoid reduces the evolution of oxygen, thereby providing a better work environment for the Rubisco enzyme.

## Conclusions

Exploring the exquisite antenna architecture of in vivo PSs is a key step for understanding the high efficiency of the light-harvesting capability of photosynthetic proteins. In the present work, we performed cryogenic single-pixel excitation-emission spectroscopy of *Chlamydomonas* cells, which enables simultaneous analysis of Chl-*b*/*a* ratio and PSII/PSI intensity ratios in the same pixel. Using this method, we investigated the LHCs that are energetically connected to PSI within the intact chloroplast.

Our results revealed the large diversity of the antenna composition of in vivo PSIs. These measurements provide evidence to answer a fundamental question as to whether the in vivo local PSII/PSI intensity ratio mainly reflects their relative stoichiometry in *Chlamydomonas* cells except for a few exceptional domains. This is crucial for studies that use fluorescence intensity to trace the structural changes of thylakoid membranes or that depend on the spatial separation of PSII and PSI. Remarkably, we found that the inhomogeneity in the antenna compositions of PSIs is related to the intracellular microscopic space. The quantitative estimation of antenna size lets us indicate the existence of mega-antenna systems of PSI in the characteristic regions surrounding the pyrenoids. Collectively, this work established an important framework based on spectroscopic techniques to estimate the in vivo antenna configuration of PSI supercomplexes that can be extended to study the antenna diversity in other organisms.

## Materials and methods

### Culture and Preparation of *Chlamydomonas* Cells

In the present experiments, a wild-type (WT) *Chlamydomonas* strain 137c (CC-125) was used. The strain was grown on agar plates containing a Tris-acetate-phosphate (TAP) medium (15 g agar powder/L TAP solution) under low white light (20 µE·m^−2^·s^−1^). These cells were transferred to a liquid TAP medium and shaken in a rotary shaker (220–230 rpm) at 23°C with the illumination of white light (30 µE·m^−2^·s^−1^) for three to four days. Subsequently, cells were cultivated photo-autotrophically in a liquid high-salt medium (HSM) under low white light (40 µE·m^−2^·s^−1^) for 12–24 hours. Before each microscope measurement, the cell suspension was sealed into a home-built sample holder and then rapidly cooled to about 80 K within 25 minutes. To avoid the occurrence of any physiological mechanisms, the sample loading and cooling processes were carried out under weak green light.

### Isolation of PSI–LHCI Supercomplexes

Thylakoid membranes from the WT were isolated as previously reported (57). To purify the photosynthetic supercomplexes from the isolated thylakoids, amphipol A8-35 was used as previously reported (58). In brief, thylakoid membranes were detergent solubilized as described in a previous report (57), and then immediately mixed with A8-35 at a final concentration of 1.0% and incubated on ice for 15 minutes. The A8-35-treated membranes were separated by detergent-free sucrose density gradient ultracentrifugation at 230,000×g for 16 hours.

### Preparation of a Trimeric PSI Complex from Cyanobacterium

For isolation of the trimeric PSI, we used the thermophilic cyanobacterium *T. vestitus* 47-H strain, in which a 6-histidine tag was introduced onto the C-terminus of the CP47 subunit in PSII. The thylakoid membranes were prepared according to the method of Nakamura et al. (59). Thylakoids were solubilized with 1% (w/v) *n*-dodecyl-β-D-maltoside (β-DDM) and then applied to a Ni^2+^ affinity column as described previously (59). Fractions that do not include PSII were collected by washing the column with a 40 mM MES-NaOH (pH 6.0) buffer containing 1.0 M betaine, 10% (w/v) glycerol, 10 mM CaCl_2_, 10 mM MgCl_2_, 100 mM NaCl, 20 mM imidazole, and 0.03% β-DDM and subsequently diluted with an equal volume of 50 mM HEPES–NaOH (pH 7.5) buffer containing 25% glycerol, 5 mM CaCl_2_, 10 mM MgCl_2_, and 0.03% β-DDM (buffer A). The diluted sample was applied to a DEAE-Toyopearl 650S (Tosoh) column equilibrated with buffer A containing 50 mM NaCl (buffer B). The column was washed with buffer B until the eluate became colorless. The PSI-trimer-enriched fraction was eluted with buffer A containing 500 mM NaCl and stored at −80°C. The purified PSI samples were diluted in a 20 mM tricine buffer (pH 7.5) containing 25 mM MgCl_2_ and 0.02% (w/v) β-DDM. Before cryo-ESM measurement, the PSI solution was centrifuged at a rotation speed of 20,000 g for 300 s at 4°C to ensure the removal of aggregated components. The concentration of the PSI solution was adjusted to 50 nM by the tricine buffer.

### Steady-state Measurements

A conventional fluorometer (F4500, Hitachi High-Tech, Tokyo, Japan) was used to carry out the steady-state fluorescence measurements. Based on previous reports, we confirmed that light induction is sufficient for the occurrence of ST mechanisms of *Chlamydomonas* (22, 23). Briefly, the ST induction light was supplied by a halogen tungsten lamp. Far-red light preferentially stimulating PSI and blue light preferentially exciting PSII were prepared with band-pass filters centered at 710 nm (ca. 178 µE·m^−2^·s^−1^) and 469 nm (ca. 120 µE·m^−2^·s^−1^), respectively. The light was also transmitted through a flat bottle containing liquid water to attenuate the thermal radiation. For the low-temperature measurement of the cells, *Chlamydomonas* cell suspensions were contained in a copper sample holder sealed with greased acrylic windows, and then immediately immersed in the liquid N_2_ in the Dewar vessel. To avoid the re-absorption effect, the optical density of the cell concentration was adjusted to ca. 0.1.

### High-speed Cryogenic Excitation Spectral Microscope (cryo-ESM)

Line-laser-type microscopes based on monochromatic or multicolor lasers have been developed and used to study photosynthetic organisms and membranes (22, 23, 33, 60). Here, we use a cryo-ESM equipping a line laser to enable the rapid acquisition of the excitation spectrum of every pixel of a microscope image (33). Recently, we updated this optical system to a low-temperature-compatible mode by integrating it into a designed cryostat (38). In the excitation pathway of cryo-ESM, a white excitation light is generated by a photonic-crystal fiber (PCF) (SCG-800, Newport, Irvine, CA, USA) pumped by a Ti:S laser at 758–765 nm. Then, the light is dispersed by a prism to form a wavelength-dispersed multi-color line focusing on the focal plane of the objective lens. Excitation light from 635 nm to 695 nm is selected by filtering through a slit and is reflected onto the objective by a DM (FF700-Diol-25×36, Semrock, Rochester, NY, USA) (Fig. 1). In the emission pathway, the fluorescence beyond the cut-on wavelength of 700 nm of the DM is injected into a polychromator (SpectraPro 320i, Acton Optics, Acton, MA, USA) and detected by an electron multiplying charge-coupled device (EMCCD) camera (Pro EM-HS-A, Teledyne Princeton Instruments, Trenton, NJ, USA). The emission spectra in the range of 700–743 nm upon different excitation wavelengths are used to reconstruct the excitation spectrum (Fig. 1A). We corrected the fluctuation in the spectral profile of the excitation laser by measuring its spectrum at the same time as that of the reference. See our previous works for more information (22, 33).

One can also acquire the emission spectra simply by replacing the DM with an HM and setting a 640 nm band-pass filter in the excitation pathway (Fig. 1B). In the emission spectral mode of cryo-ESM, we can measure the 670–750 nm emission spectrum of *Chlamydomonas* cells at low temperatures. The wavelength dependence of the sensitivity of the system was estimated by a spectral measurement of a photometric standard lamp (JPD100V500WCS, Ushio Lighting, INC, Tokyo, Japan). Figure S15 shows the measured emission spectrum and the calculated emission spectrum based on Planck’s law in the 665–751 nm regions. Planck’s curve is given by a formula below (61):

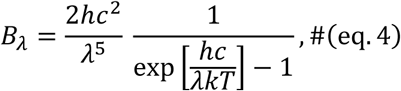

where *h* and *k* are the Planck and Boltzmann constants, respectively, and *c* is the speed of light. λ is the wavelength. The calibration curve was estimated by dividing the obtained spectrum by the theoretical curve in eq. 4. It was used to correct the emission spectrum obtained by the microscopy experiment.

All measurement procedures and data analyses were conducted via LabVIEW (National Instruments, Austin, TX, USA), Python, and Igor Pro 9 (WaveMetrics, Inc., Portland, OR, USA) software.

## Acknowledgments

We thank Professor Jun Minagawa (National Institute for Basic Biology), Professor Ryutaro Tokutsu (Kyoto University), and Professor Eunchul Kim (National Institute for Basic Biology) for the kind gift of the *Chlamydomonas* cell strain and the isolated PSI–LHCI supercomplex. We also thank Professor Takumi Noguchi (Nogao University) and Dr. Ryo Nagao (Shizuoka University) for the gift of the PSI trimer isolated from cyanobacteria.

## Author contributions

Y.S. and X.Z. conceived and designed the study. R.T. performed the experiment of the isolated sample that was indispensable to estimate the in-vivo PSI antenna size. X.Z. performed the other experiments and data analysis. Y.S. and X.Z. wrote the manuscript with contributions from all authors. S.Y. supervised the experiments and writing process. All authors read and approved the final manuscript.

